# SpyRing-Mediated Cyclization of TEV Protease, Guided by AlphaFold, Improves Thermostability

**DOI:** 10.1101/2025.03.30.646106

**Authors:** Tadashi Nakai, Yota Nakai, Naoki Takami, Emi Nakai, Toshihide Okajima

**Affiliations:** Graduate School of Science and Technology, Hiroshima Institute of Technology, Hiroshima 731-5193, Japan; Faculty of Life Sciences, Hiroshima Institute of Technology, Hiroshima 731-5193, Japan; Hiroshima University Senior High School, Hiroshima 734-0005, Japan; Institute of Scientific and Industrial Research (SANKEN), Osaka University, Ibaraki, Osaka 567-0047, Japan

**Keywords:** TEV protease, SpyRing-mediated cyclization, Protein stabilization, SpyTag/SpyCatcher, AlphaFold design, Synthetic biology

## Abstract

Cyclization is a promising strategy to enhance protein stability, but its applicability is often limited by structural constraints such as the distance between terminal regions. Here, we report the rational design and characterization of a cyclized Tobacco Etch Virus protease (cTEVp) using the SpyRing system, which enables covalent cyclization through SpyTag/SpyCatcher-mediated isopeptide bond formation. We applied this approach to a widely used engineered TEVp variant (L56V, S135G, S219V, Δ238–242) and employed AlphaFold structure prediction to optimize linker length and domain positioning. Despite the ~40 Å separation between the N- and C-termini of native TEVp, AlphaFold modeling suggested that the fused SpyTag and SpyCatcher domains can adopt a favorable configuration for intramolecular cyclization. The resulting cTEVp exhibited proteolytic activity comparable to the non-cyclized TEVp, indicating that structural constraint via SpyRing-mediated cyclization did not impair enzymatic function. Importantly, cTEVp displayed significantly improved thermostability relative to its non-cyclized counterpart, as demonstrated by higher retention of soluble enzyme and residual activity following heat treatment at 50°C. Our findings validate the effectiveness of SpyRing-mediated cyclization in improving TEVp stability and highlight the utility of computationally guided cyclization as a generalizable strategy for engineering thermally resilient proteins. This study establishes a framework that integrates structure prediction and rational protein engineering, contributing to the development of robust biocatalysts for synthetic biology and industrial biotechnology applications.

**Table of Contents (TOC) / Abstract Graphic:** 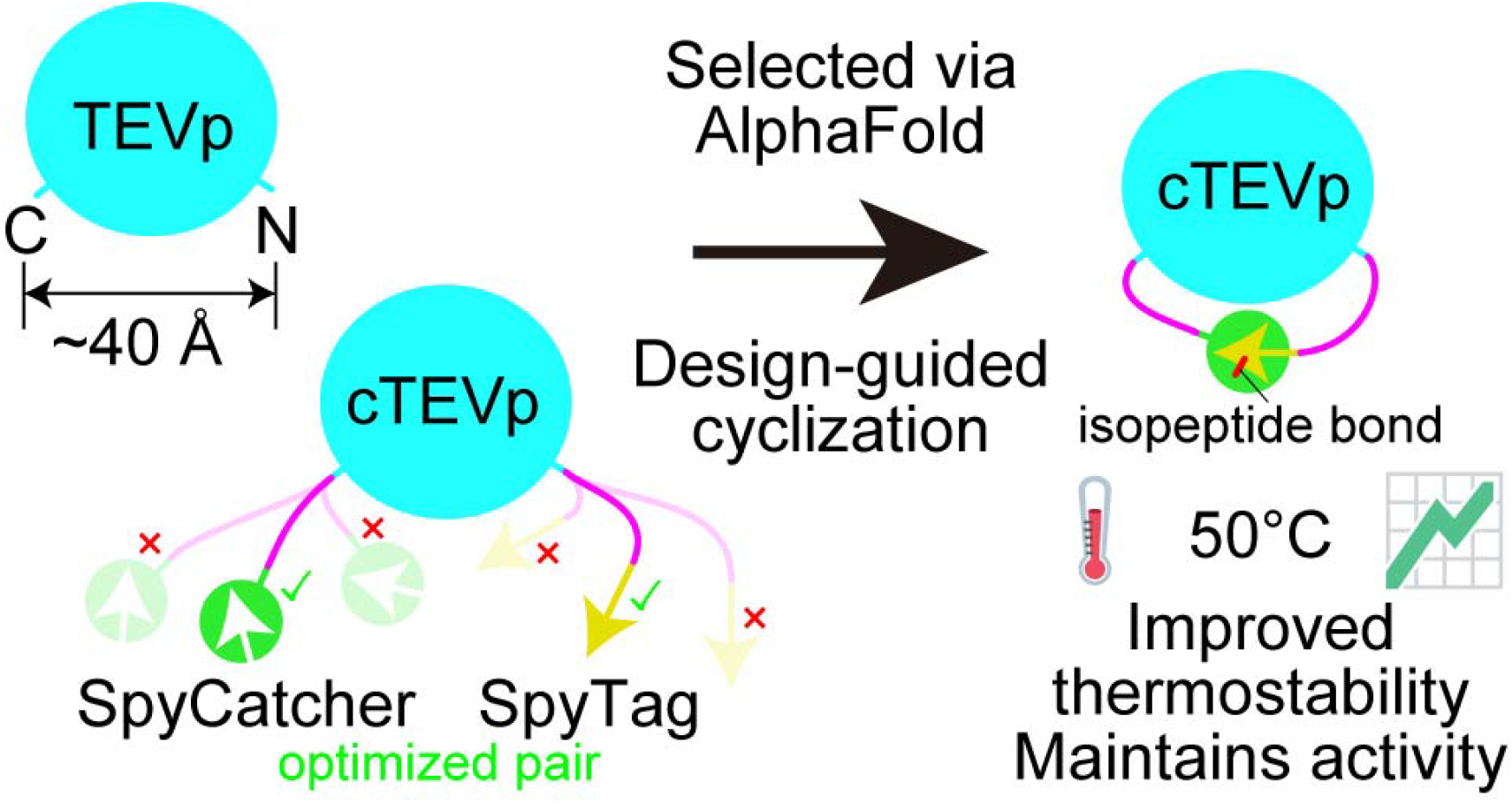

## 1. Introduction

Recent advances in biotechnology and bioengineering have dramatically improved the efficiency of recombinant protein production. In particular, the rise of synthetic biology has enabled modular and rational design of biomolecular systems, allowing for efficient production and engineering of recombinant proteins. In parallel, the addition of peptide tags or fusion partners to target proteins has become a standard approach for construct design, enabling efficient purification by affinity chromatography.^1^ To obtain recombinant proteins in a near-native form, it is necessary to remove these tags using specific proteases. Among them, the Tobacco Etch Virus protease (TEVp), which exhibits high substrate specificity, is commonly employed for tag removal.^2,3^ However, wild-type TEVp suffers from poor solubility, autolysis, and low yield, which have long hindered its production and utility.

To address these issues, numerous engineering strategies have been explored for TEVp. For example, substitution at the S219 residue to suppress autolysis has been investigated, and the S219V mutant has been reported to exhibit approximately 100-fold higher stability compared to the wild-type enzyme.^4^ Additionally, directed evolution experiments have shown that introducing three additional mutations (T17S, N68D, and I77V) into the parental S219N mutant enhances protein yield by approximately fivefold.^5^ Furthermore, a C-terminal deletion mutant lacking residues 238–242 (Δ238–242) has been demonstrated to improve solubility while retaining catalytic activity comparable to the wild-type.^6^ Other studies have reported that the L56V and S135G mutations further enhance solubility,^7,8^ and that combining these mutations (L56V, S135G, S219V, and Δ238–242) enables large-scale production of TEVp.^9^ Moreover, incorporating additional mutations (T17S, N68D, and I77V) into this background has been shown to improve solubility, stability, and catalytic activity, leading to a substantial increase in yield.^10^ More recently, AI-assisted rational design has been employed to generate next-generation TEVp variants with enhanced expression, stability, and catalytic activity.^11^ Although this extensive engineering, which involves substitutions at more than 15% of the total amino acid residues, has significantly improved catalytic activity and thermostability, these variants have only been used in limited applications and require further evaluation for enzymatic properties including stringent substrate specificity.

In this study, we aimed to further enhance the stability of TEVp by introducing a cyclization strategy using the SpyRing method into the TEVp variant containing the L56V, S135G, S219V mutations and the C-terminal deletion Δ238–242,^9^ which is commonly used due to its improved solubility and activity. Here, we refer to this variant as the standard TEVp. The SpyRing technique employs the self-assembling SpyTag (ST) and SpyCatcher (SC) system, consisting of a peptide and a small protein domain,^12^ widely used as orthogonal biological parts in synthetic biology. By genetically fusing these elements to the N- and C-termini of a target protein, we enable the spontaneous and irreversible formation of an intramolecular covalent bond, resulting in protein cyclization.^13^ While previous reports have suggested that SpyRing-mediated cyclization may not be suitable for proteins with N- and C-terminal distances exceeding 15 Å,^12^ we selected TEVp as a challenging model system, given that its termini are separated by approximately 40 Å in the native structure.^14^ Structural models of cyclized TEVp (cTEVp) generated using the SpyRing system are shown in Figure 1. The major advantage of this approach is that covalent cyclization helps preserve the overall topology of the protein, thereby promoting efficient refolding and recovery of activity after thermal stress.^13,15^ Moreover, this system is highly convenient compared to conventional methods for the cyclization, as it does not require additional reagents or enzymes and functions both *in vivo* and *in vitro*. For example, significant improvements in thermostability have been reported when the SpyRing system was applied to various enzymes, including β-lactamase,^13^ phytase,^15^ xylanase,^16^ and luciferase.^17^ Notably, several of these enzymes, such as phytase and xylanase, are widely used in industrial applications, including animal feed supplementation and biomass degradation processes. Beyond industrial applications, the SpyTag/SpyCatcher system has also been explored in diverse biomedical applications,^18^ including vaccine design, drug delivery systems, and cell surface display technologies, underscoring its versatility across both industrial and therapeutic fields.

**Figure 1.**
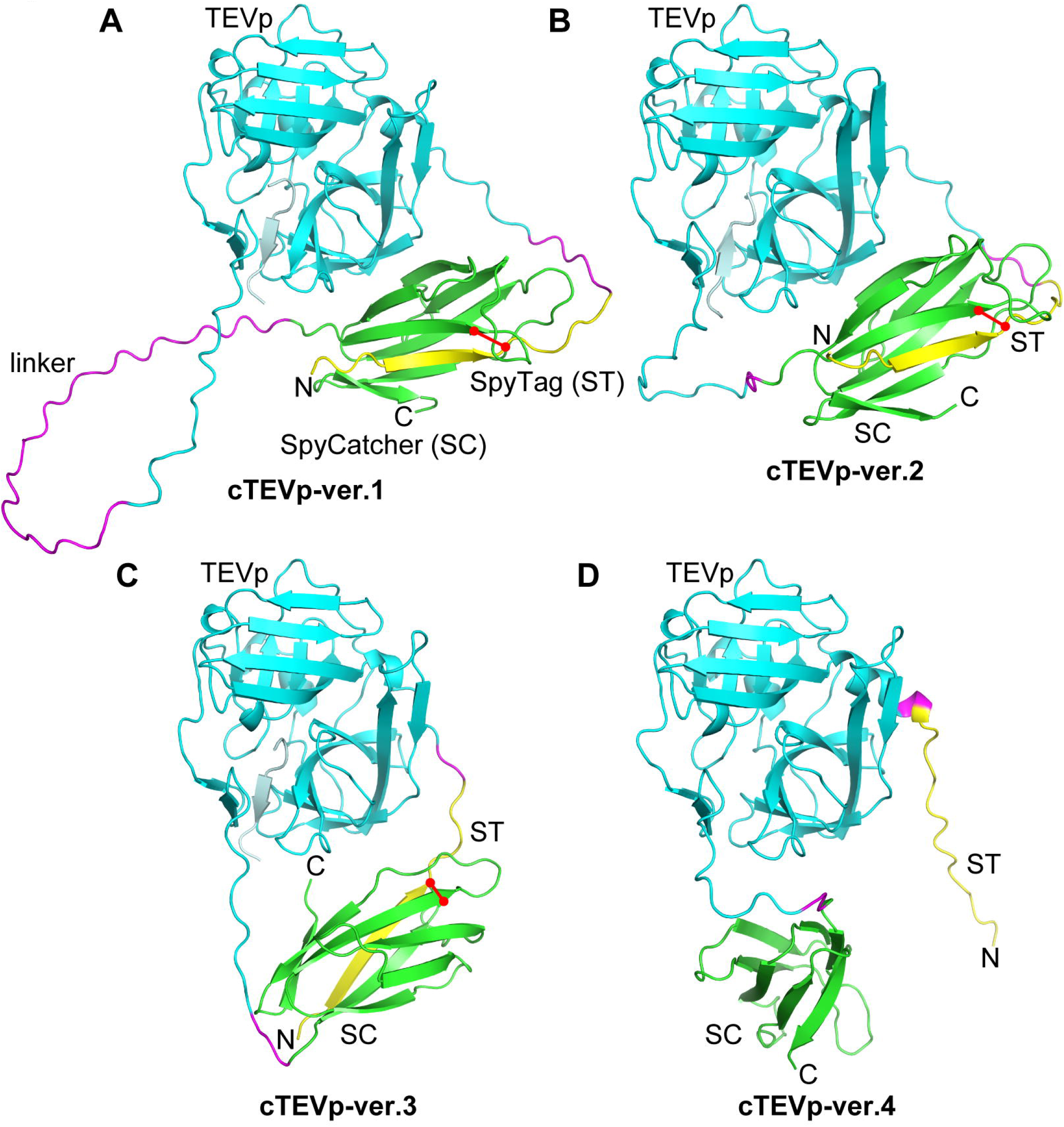
Structural models of various cTEVp designs generated using AlphaFold. Representative models from multiple linker variants are shown. **(A)** The initial model constructed by directly attaching the ST, SC, and linker sequences reported in ref.^12^ to the N- and C-termini of TEVp. **(B)** A model generated by removing 30 amino acid residues from the linker region of ver.1. This model was selected as the final design used in this study. **(C)** A model generated by further removing 17 amino acid residues from ver.2 to further shorten the linker region. **(D)** A model in which one additional residue was removed from ver.3. In this model, cyclization was structurally disrupted, preventing ST and SC from forming a bond. TEVp is shown in cyan, SpyCatcher (SC) in green, SpyTag (ST) in yellow, and the linker region in magenta. The TEVp substrate (GENLYFQG) is displayed in pale cyan, and the isopeptide bond formed between SC and ST is highlighted in red. N-terminal His-tags and C-terminal Strep-tags are not included in these models.

In this paper, we constructed cTEVp by applying the SpyRing method to the standard TEVp (L56V, S135G, S219V, Δ238–242), a widely used conventional mutant with improved solubility and activity,^9^ and evaluated its biochemical properties. Leveraging AlphaFold structure prediction,^19^ we rationally designed a cTEVp by optimizing the spatial arrangement of SpyTag and SpyCatcher fused to the termini, despite the native ~40 Å separation between the N- and C-termini. We demonstrate that cTEVp retains proteolytic activity comparable to that of the standard TEVp while exhibiting improved thermostability. These results validate the applicability of the SpyRing method to TEVp and showcase a design-driven approach that integrates computational modeling with protein engineering, offering a versatile strategy for enhancing protein stability in synthetic biology.

## 2. Results

### 2.1. Structural design and optimization of cTEVp

Recent advances in artificial intelligence (AI) technology have enabled highly accurate prediction of protein three-dimensional structures based on amino acid sequences. In this study, we utilized AlphaFold 3^19^ to optimize the design of cTEVp using the SpyRing system. Initially, we constructed a model (cTEVp-ver.1) by attaching the ST, SC, and linker sequences reported in ref.^12^ directly to the N- and C-termini of TEVp (Figures 1A and S1A). In this model, the 30-residue linker between TEVp and SC appeared excessively flexible in the AlphaFold-predicted structure, with the corresponding region exhibiting very low pLDDT scores (pLDDT < 50), indicating reduced confidence in its structural order (Figure S2A). These features suggest that such a linker length may not be ideal for effective stabilization via cyclization.

Therefore, we iteratively revised the design by shortening and adjusting the linker region based on the AlphaFold-predicted structures. While multiple variants with different linker lengths and compositions were evaluated during the optimization process, representative models are shown in Figure 1. Specifically, ver.2 was generated by removing 30 residues from the linker of ver.1 (Figure 1B and S1B). In this model, SpyTag and SpyCatcher were brought into proximity and formed a complex, while maintaining sufficient flexibility and spacing from the TEVp termini to permit cyclization. This configuration was predicted to be the most structurally favorable among the tested designs. To achieve further compaction, ver.3 was designed by removing an additional 17 residues from ver.2 (Figure 1C and S1C). While intramolecular cyclization was still predicted by AlphaFold, the overall configuration appeared more constrained, with reduced spacing between TEVp and the fused domains. Although Ramachandran analysis did not reveal significant backbone strain, the shorter linker may have reduced the flexibility at the termini, potentially affecting proper folding or positioning for cyclization. Experimental evaluation of ver.3 supported this interpretation: the purified protein exhibited reduced cyclization efficiency, increased oligomer formation, and decreased thermostability compared to ver.2 (data not shown). Finally, in ver.4 (Figure 1D and S1D), where one more residue was removed from ver.3, the linker became too short, and cyclization was structurally impeded.

Based on these results, ver.2 was selected as the most suitable design in terms of both structural feasibility and predicted stability, and all subsequent experiments were performed using this construct. Notably, AlphaFold modeling indicated that although the N- and C-termini of TEVp are separated by approximately 40 Å in the native structure, the fused SpyTag and SpyCatcher domains were positioned favorably, enabling successful intramolecular cyclization (Figure 1B).

### 2.2. Protein expression and purification

In this study, we constructed a new high-copy-number expression vector, pHIT184 (Figure S3), by combining the expression cassette of a T7 promoter-based system with a pUC-derived replication origin. This vector, which retains the IPTG-inducible T7 promoter and *lacI*/*lacO* regulatory elements, is advantageous for genetic manipulation and enables high-level protein expression of the inserted gene in *Escherichia coli*. We subsequently evaluated its utility by expressing several TEVp variants and substrate proteins, obtaining sufficient amounts of recombinant proteins from small-scale (100 mL) cultures. Our system yielded 94 mg/L of purified protein, which is higher than the previously reported yield of TEVp (50 mg/L).^9^ Since the amino acid sequence of the enzymatic region is identical in both proteins (except for the addition of 11 residues, including a C-terminal Strep-tag, in our construct), this result confirms the high efficiency of our expression and purification system. Notably, both the TEVp and substrate genes used in this study were synthesized as codon-optimized sequences for *E. coli*, achieving high expression without the need for additional rare codon tRNA supply plasmids (e.g., pRIL). Furthermore, TEVp produced by this system has been routinely used in our laboratory for the removal of affinity tags from various recombinant proteins without any issues, demonstrating its practical applicability.^20^

In addition to the standard TEVp described above, two other TEVp variants—cTEVp and ncTEVp—were similarly expressed in *E. coli* and purified to high purity using affinity chromatography. The purified preparations of all three TEVp variants are shown in Figure 2A (lanes 3, 7, and 11). Using the same method, we also purified three substrate proteins: a monomeric variant of the T7 phage Ocr protein (Mocr),^21^ T4 lysozyme (T4L),^22^ and chitin-binding domain (ChBD).^23^ T4L is a well-established fusion partner widely used in structural biology, whereas Mocr has been reported to enhance the solubility of fusion proteins and is compatible with structural studies. ChBD, derived from a thermostable archaeal chitinase, offers high thermal resistance suitable for heat stress assays. Details of all purification results are summarized in Table 1. SDS-PAGE analysis (Figure 2A) showed that the major bands of TEVp and ncTEVp corresponded well with their theoretical molecular weights (Table 1). In contrast, the cTEVp band shifted to a lower molecular weight compared to ncTEVp, although the only difference between them is a single residue substitution (Figure 2A, lane 7). This shift is likely due to altered electrophoretic mobility caused by the cyclized structure, as has been reported for other SpyRing enzymes.^13,15,16^ Moreover, several ladder bands were detected at higher-molecular-weight positions for cTEVp, likely due to intermolecular SpyTag-SpyCatcher cross-linking, a phenomenon commonly seen with other SpyRing enzymes.^13,15,16^

**Table 1.** Recombinant proteins expressed in this study.

| **Protein Name** | **Expression Temperature (°C)** | **Induction Time (hours)** | **Yield (mg) <sup>a</sup>** | **Theoretical Molecular Weight <sup>c</sup>** | **Reference** |
| --- | --- | --- | --- | --- | --- |
| TEVp | 25 | 28 | 9.4 <sup>b</sup> | 29061.89 | This study <sup>d</sup> |
| cTEVp | 25 | 24 | 3.8 <sup>b</sup> | 39857.95 | This study |
| ncTEVp | 25 | 24 | 0.8 <sup>b</sup> | 39813.94 | This study |
| Mocr | 37 | 16 | 9.2 | 18141.49 | 21 |
| T4L | 25 | 18 | 5.7 | 22495.44 | 22 |
| ChBD | 37 | 16 | 16.6 | 16459.96 | 23 |

<sup>a</sup> Yield per 100 mL of culture medium.
<sup>b</sup> TEVp, cTEVp, and ncTEVp were expressed as MBP fusion proteins and cleaved intracellularly by their own TEVp activity; the values shown here are calculated after MBP removal.
<sup>c</sup> Theoretical molecular weight was calculated using the Compute pI/Mw tool available at Expasy (https://web.expasy.org/compute_pi/).
<sup>d</sup> Differences between this study and ref. <sup>9</sup> are shown in Table S3.

**Figure 2.**
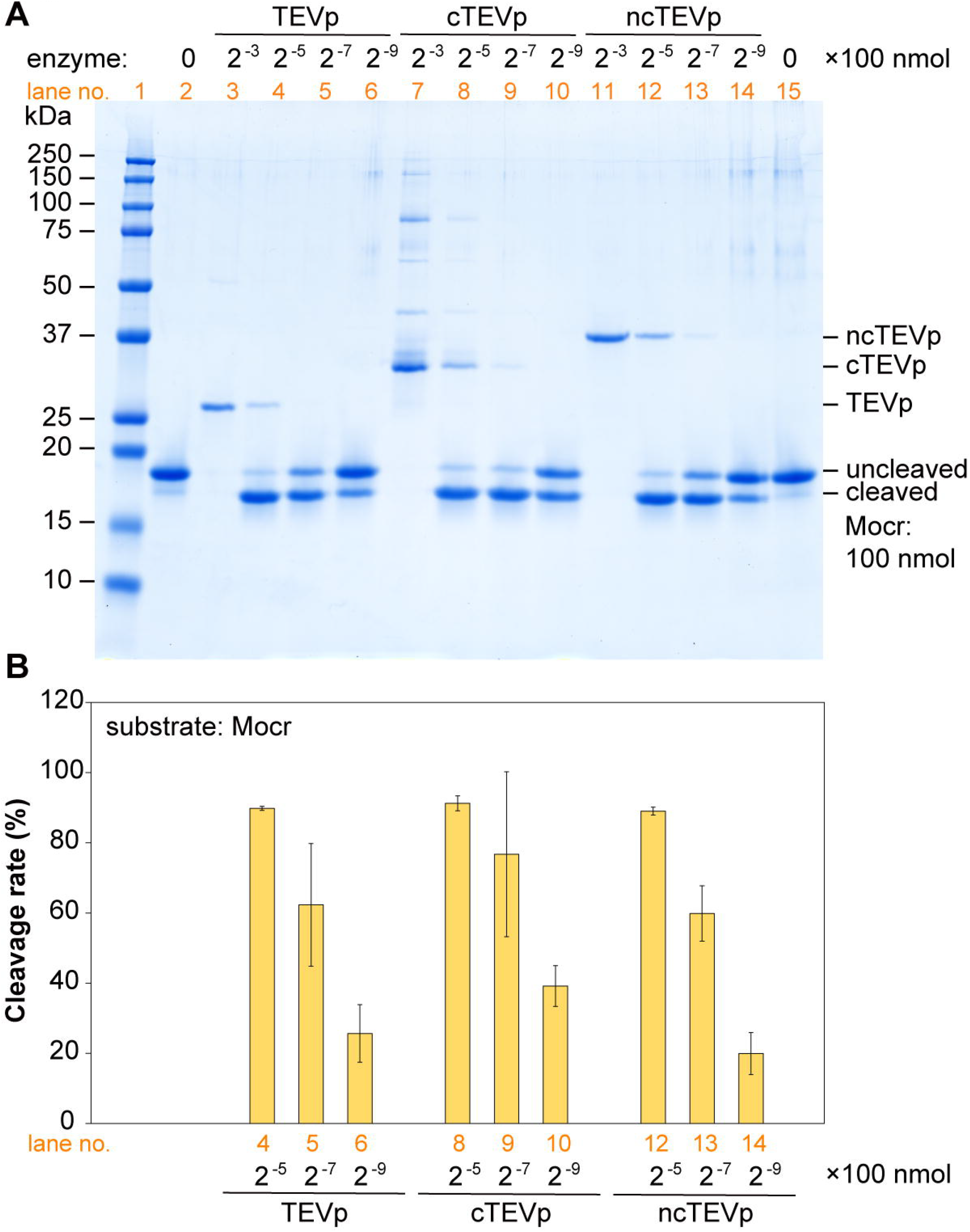
SDS-PAGE analysis of purified enzymes and substrate proteins, and quantitative analysis of substrate cleavage activity. **(A)** SDS-PAGE analysis using Mocr as the substrate. Lane 1: Molecular weight markers. Lanes 2 and 15: substrate only control (Mocr, 100 nmol); Lanes 3, 7, and 11: enzyme only (100 × 2^−3^ nmol), without substrate; Lanes 4, 8, and 12 enzyme (100 × 2^−5^ nmol) + substrate (100 nmol); Lanes 5, 9, and 13: enzyme (100 × 2^−7^ nmol) + substrate (100 nmol); Lanes 6, 10 and 14: enzyme (100 × 2^−9^ nmol) + substrate (100 nmol). Enzyme and substrate were mixed and incubated at 37°C for 60 minutes prior to electrophoresis. **(B)** Quantitative analysis of substrate cleavage rates. Cleavage rates (%) were calculated based on the integrated intensity of the uncleaved Mocr bands. The initial band intensity (uncleaved substrate) was set to 100%, and cleavage rate was determined by subtracting the intensity of remaining substrate after the reaction. Data represent mean ± SD from three independent experiments.

Initially, cTEVp was expressed by fusing only SpyTag and SpyCatcher to its termini; however, the expression level was extremely low, making it difficult to obtain sufficient amounts of purified protein (data not shown). Therefore, similar to TEVp, maltose-binding protein (MBP) was fused to the N-terminus of cTEVp, which significantly improved its expression level. Subsequently, using this MBP-fused cTEVp as a template, we introduced the Asp-to-Ala mutation to generate the MBP-fused ncTEVp. Both enzymes underwent autocleavage in *E. coli*, resulting in the removal of the MBP fusion, and were ultimately isolated in a cleaved form. These results confirmed that both cTEVp and ncTEVp, like the standard TEVp, efficiently cleave large fusion partners, such as MBP, *in vivo*.

### 2.3. Evaluation of substrate cleavage activity of cTEVp

To evaluate the enzymatic activity of cTEVp, we employed three model substrates—Mocr, T4L, and ChBD—each containing a TEVp cleavage site (TEVcs) between an N-terminal H_6_-tag and the substrate protein. These substrates, described in the previous section, enabled comparative analysis of proteolytic activity under standard and heat stress conditions.

First, we compared the substrate cleavage activity of cTEVp with that of ncTEVp and the standard TEVp using Mocr as the substrate (Figure 2). SDS-PAGE analysis (Figure 2A) was performed to assess the cleavage reaction after incubating enzyme and substrate mixtures at various enzyme-to-substrate molar ratios (1:32 to 1:512) at 37°C for 1 hour. Under a 1:128 enzyme-to-substrate molar ratio, which approximates the commonly used standard reaction condition for TEVp (1:100), all three enzymes cleaved more than 50% of the substrate (corresponding to lanes 5, 9, and 13).

Quantitative analysis of the substrate cleavage rates (Figure 2B) showed that cTEVp, TEVp, and ncTEVp all exhibited high cleavage activities. At a 1:128 molar ratio, cTEVp, TEVp, and ncTEVp achieved cleavage rates of 77 ± 24%, 62 ± 18%, and 60 ± 8%, respectively. No statistically significant differences were observed among the three enzymes (*p* = 0.44 and *p* = 0.30, both compared to cTEVp).

These results demonstrate that cTEVp, cyclized via the SpyRing system, exhibits comparable substrate cleavage activity to both TEVp and ncTEVp. Thus, despite the structural constraint imposed by cyclization, cTEVp retained practical cleavage activity equivalent to that of the standard TEVp.

### 2.4. Effect of SpyRing-mediated cyclization on the thermostability of TEVp

To assess thermostability, we used T4 lysozyme (T4L) as a model substrate, which allowed simultaneous visualization of both enzyme and substrate on SDS-PAGE due to its relatively slow cleavage kinetics. This enabled quantitative evaluation of both the soluble enzyme fraction and residual protease activity after heat treatment.

To evaluate the thermostability of TEVp, ncTEVp, and cTEVp, we analyzed the amount of enzyme remaining in the soluble fraction and the residual protease activity after heat treatment at 50°C for 0 to 32 minutes (Figure 3). Upon heat treatment, TEVp and ncTEVp rapidly lost their soluble fraction, with almost no detectable soluble protein remaining after 8 minutes (Figure 3B). In contrast, cTEVp retained a higher proportion of its soluble fraction, with 31.6% remaining after 8 minutes, compared to 4.6% and 2.3% for TEVp and ncTEVp, respectively. These differences were statistically significant (*p* = 0.015 and *p* = 0.021, respectively, relative to cTEVp).

**Figure 3.**
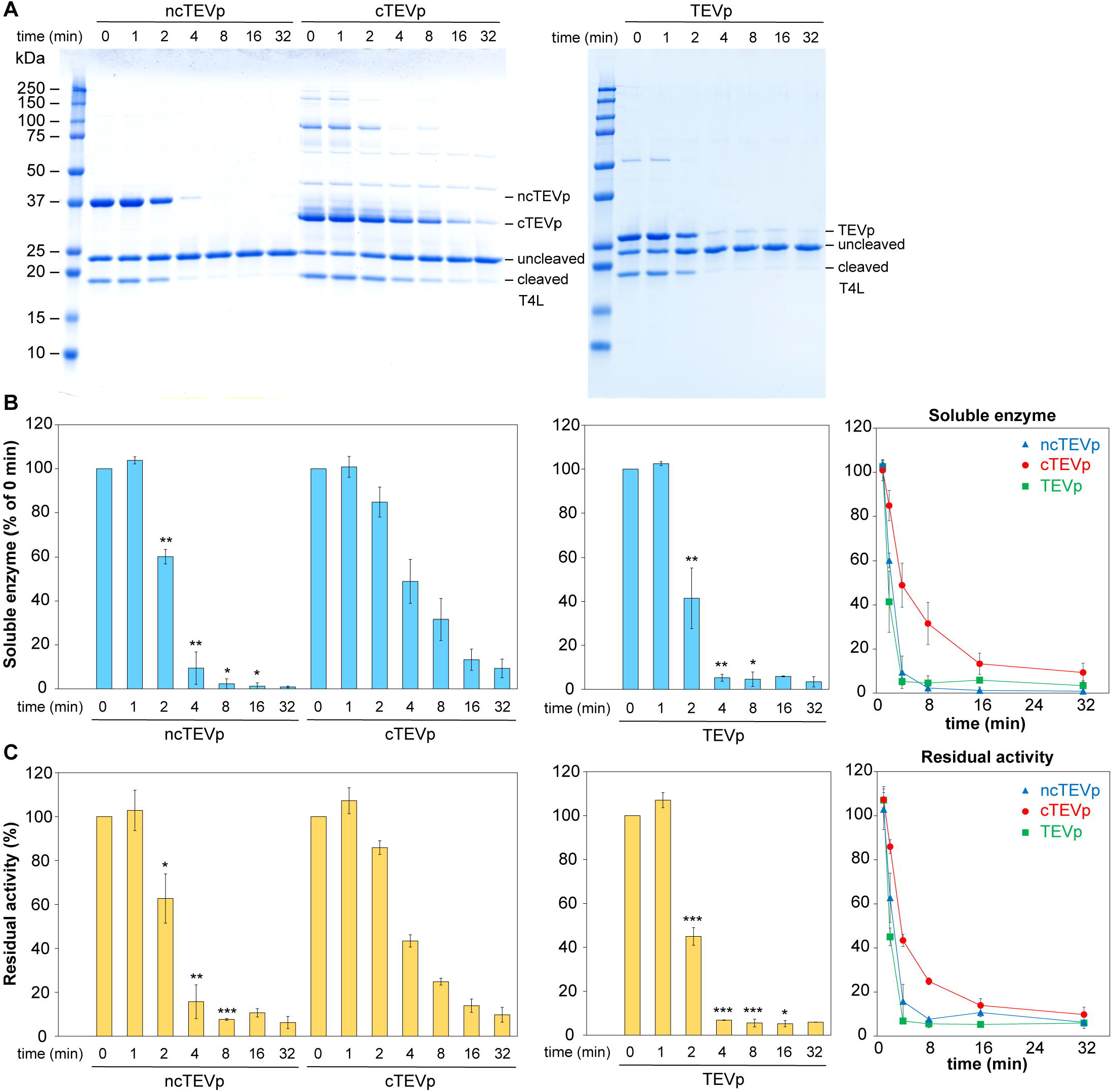
Thermostability of TEVp variants: time-course analysis of soluble fraction and residual activity after heat treatment. **(A)** SDS-PAGE analysis of soluble enzyme fractions and substrate cleavage products. Enzymes (ncTEVp, cTEVp, and TEVp) were incubated at 50°C for the indicated durations (0, 1, 2, 4, 8, 16, or 32 minutes), followed by centrifugation to separate soluble fractions. These soluble fractions were incubated with equimolar amounts of T4L substrate at 37°C for 3 hours. The reaction mixtures were analyzed by SDS-PAGE to assess the amount of soluble enzyme and residual enzymatic activity. Bands corresponding to enzyme variants (ncTEVp, cTEVp, TEVp), uncleaved and cleaved T4L are indicated. **(B)** Quantification of soluble enzyme fraction after heat treatment. Bar graphs and line plots show the percentage of soluble enzyme remaining in the soluble fraction after incubation at 50°C for the indicated durations. Band intensities were quantified from SDS-PAGE images in (A), with the 0 min time point set to 100%. **(C)** Quantification of residual enzymatic activity. Residual activity was assessed by measuring cleavage of T4L substrate by the heat-treated soluble enzyme fractions. Product band intensities were quantified and normalized to the 0 min control (set as 100%). Data represent means ± SD from multiple independent experiments (*n* = 4 for cTEVp, *n* = 2 for ncTEVp and TEVp). Asterisks indicate statistically significant differences relative to cTEVp at the same time point (\**p* < 0.05, \*\**p* < 0.01, \*\*\**p* < 0.001; see Methods for details).

We next evaluated the proteolytic activity of the heat-treated soluble enzyme fractions using a substrate cleavage assay (Figure 3C). The time-dependent decline in residual activity correlated well with the remaining soluble fraction, and cTEVp consistently retained substantially higher activity throughout the time course. At multiple time points, particularly at 8 minutes and beyond, the residual activity of cTEVp was significantly higher than that of TEVp and ncTEVp. For example, after 8 minutes at 50°C, cTEVp retained 24.8% of its activity, whereas TEVp and ncTEVp retained only 5.6% and 7.6%, respectively (*p* = 0.00016 and *p* = 0.00013).

These results demonstrate that while TEVp and ncTEVp exhibit similar thermostability profiles, the SpyRing-mediated cyclization of TEVp consistently enhanced resistance to heat-induced loss of soluble protein and enzymatic activity across multiple time points. The pronounced differences observed particularly at later time points underscore the effectiveness of covalent cyclization in maintaining enzymatic function under thermal stress.

### 2.5. Activity profiling of cTEVp at moderate and elevated temperatures

To evaluate the impact of SpyRing-mediated cyclization on the thermal robustness of TEVp activity, we first compared the catalytic efficiencies of TEVp, cTEVp, and ncTEVp using T4 lysozyme (T4L) as a standard substrate at 30°C and 37°C. While previous reports have suggested that TEVp exhibits optimal activity at 30°C,^24^ our results showed that all three variants maintained comparable activity at both temperatures (Figure 4A, left panel). Based on these results, enzyme amounts were titrated at 30°C and 37°C for subsequent comparisons under heat stress conditions.

**Figure 4.**
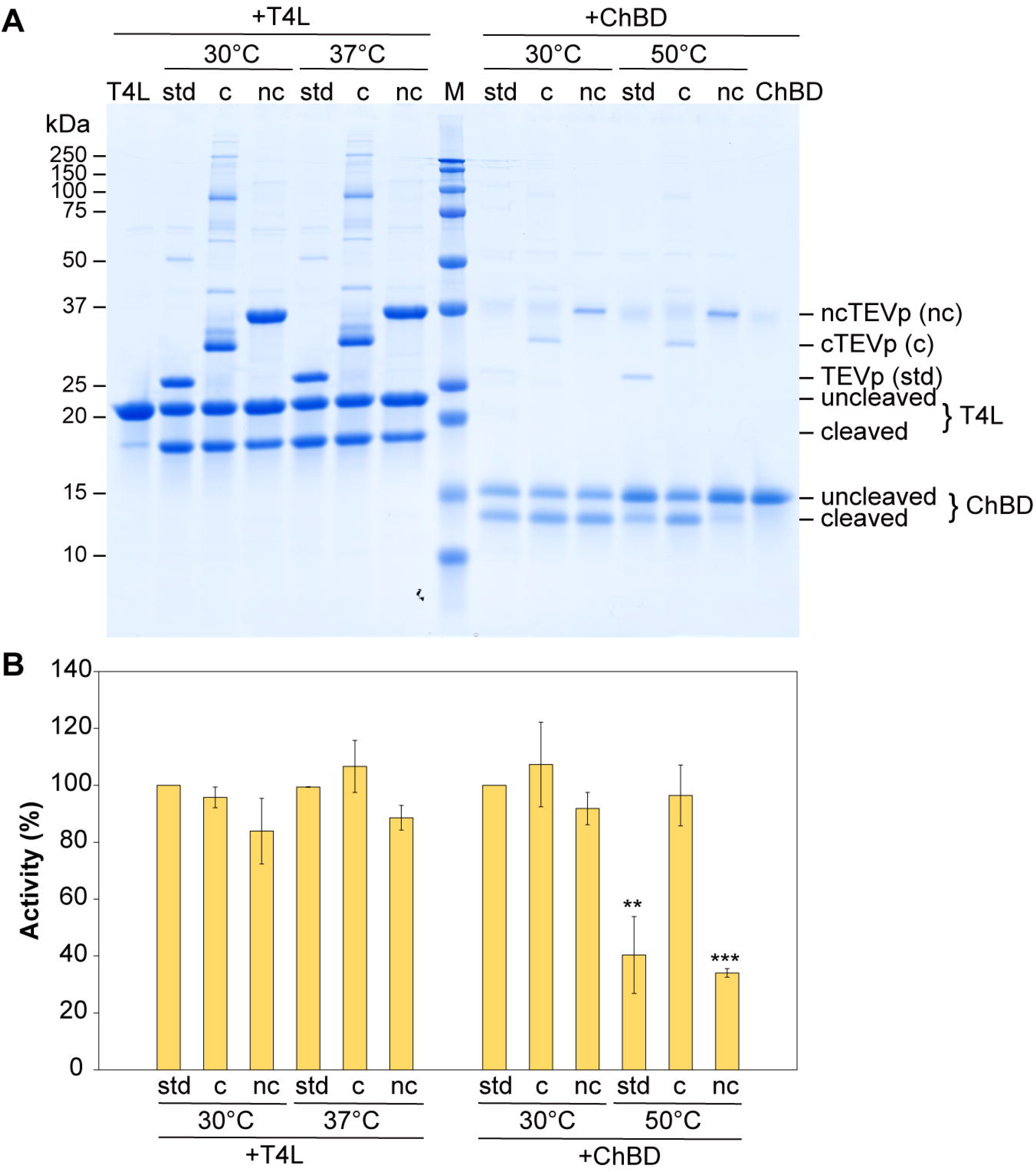
Comparative analysis of TEVp variants for proteolytic activity at moderate and elevated temperatures. **(A)** SDS-PAGE analysis of substrate cleavage by TEVp (std), cTEVp (c), and ncTEVp (nc) using T4 lysozyme (T4L) and chitin-binding domain (ChBD) as substrates. For the T4L assays (left panel), enzymes and substrate were incubated at 30°C or 37°C for 3 hours. All three TEVp variants exhibited comparable cleavage activity at both temperatures, supporting their use in normalized comparisons. For the ChBD assays (right panel), cleavage reactions were conducted at 30°C or 50°C using enzyme amounts adjusted based on the T4L results. At 50°C, cTEVp showed visibly stronger cleavage product bands than either TEVp (std) or ncTEVp, indicating superior retention of catalytic activity under heat stress. Substrate-only controls are shown at the far left (T4L) and far right (ChBD). Molecular weight markers (M) are included in the center. **(B)** Quantification of substrate cleavage efficiency from (A). For T4L (left graph), the intensity of the cleaved band from the 30°C TEVp condition was set to 100%, and all other values were normalized accordingly. For ChBD (right graph), the cleaved band intensity from the 30°C TEVp condition was likewise set as 100%. Data represent mean ± SD from three independent experiments. At 50°C, proteolytic activities of TEVp and ncTEVp were significantly lower than that of cTEVp (*p* < 0.01 and *p* < 0.001, respectively; unpaired two-tailed Student’s t-test vs. cTEVp).

Next, to assess enzymatic activity under heat stress, the same titrated enzyme amounts (adjusted based on T4L cleavage) were applied to cleavage reactions using the thermostable ChBD substrate at 50°C for 30 minutes. Because T4L proved unstable at 50°C, the thermostable ChBD substrate was used instead for comparative analysis under heat stress conditions.

As shown in Figure 4A (right panel) and quantitatively summarized in Figure 4B, cTEVp retained significantly higher proteolytic activity at 50°C than both TEVp and ncTEVp, with *p*-values of 0.0049 and 0.00056, respectively. These data demonstrate that SpyRing-mediated cyclization confers enhanced thermal resilience to the catalytic function of TEVp, even under conditions that substantially reduce the activity of its non-cyclized counterparts.

## 3. Discussion

In this study, we successfully generated a cTEVp using the SpyRing system and characterized its properties. The cTEVp retained substrate cleavage activity comparable to both ncTEVp and standard TEVp (Figure 2), demonstrating that structural stabilization via SpyTag-SpyCatcher-mediated cyclization did not adversely affect the enzymatic activity.

In contrast, comparative analysis of thermostability revealed that cTEVp consistently retained a higher proportion of soluble enzyme and residual activity than both TEVp and ncTEVp throughout the heat treatment (Figure 3). These findings indicate that cyclization via the SpyRing system stabilizes the molecular structure of TEVp and confers resistance to heat-induced inactivation. The observed loss of soluble enzyme and activity in TEVp and ncTEVp upon heat treatment is likely due to heat-induced aggregation, although the contribution of partial denaturation cannot be entirely excluded.

Although ncTEVp retains the ST and SC domains at its N- and C-termini, it was designed to prevent isopeptide bond formation by introducing a point mutation in SpyTag, thereby eliminating covalent cyclization. Structural prediction using AlphaFold suggested that ST and SC remain associated in ncTEVp despite the absence of isopeptide bond formation, indicating that cTEVp and ncTEVp are structurally highly similar. Furthermore, TEVp and ncTEVp exhibited comparable behaviors in terms of their remaining soluble fractions and proteolytic activities, both serving as indicators of thermostability. These observations suggest that the cyclized structure itself— specifically, the isopeptide bond between SpyTag and SpyCatcher—is the primary factor contributing to the enhanced stability of TEVp. This finding aligns with previous reports showing that SpyRing-mediated cyclization enhances the thermal resilience of other model enzymes, including β-lactamase^13^ and phytase.^15^

Notably, the TEVp variant used in this study already incorporates the L56V and S135G mutations found in the TEVp^2M^ construct, which has been reported to improve thermal tolerance above 40°C.^8^ Despite the presence of these stabilizing mutations, our non-cyclized variants (TEVp and ncTEVp) still showed substantial aggregation and loss of activity upon incubation at 50°C, indicating that these mutations alone are insufficient to fully protect the enzyme under thermal stress. This underscores the additive effect of SpyRing-mediated cyclization in further enhancing thermal resilience, beyond what is achievable through conventional mutagenesis. The improved performance of cTEVp is likely attributable to suppression of heat-induced aggregation, rather than promotion of reversible unfolding. This interpretation is supported by the observation that both TEVp and ncTEVp rapidly lost solubility and activity upon heat treatment, consistent with irreversible aggregation. In contrast, cTEVp retained a significantly higher proportion of soluble protein and residual activity, suggesting that SpyRing-mediated cyclization prevents the onset of misfolding or aggregation, rather than facilitating refolding after denaturation. TEVp is known to lose activity irreversibly once aggregation occurs,^25^ in contrast to some other enzymes such as β-lactamase, in which cyclization confers the ability to refold after thermal denaturation.^13^ These differences highlight that the primary benefit of SpyRing-mediated cyclization in the case of TEVp lies in its ability to prevent irreversible inactivation via aggregation.

According to the crystal structure of TEVp^14^ (PDB entry: 1LVB), the distance between the N- and C-termini is approximately 40 Å, which is relatively long. Interestingly, despite this separation, AlphaFold modeling indicated that the fused SpyTag and SpyCatcher domains were positioned favorably for intramolecular cyclization. This suggests that the flexibility and orientation of these domains can compensate for the relatively long native terminal distance, facilitating successful cyclization. Although it has been suggested that proteins with terminal distances exceeding 15 Å may not be suitable for SpyRing-mediated cyclization,^12^ the present findings challenge this assumption. Indeed, while phytase (with a terminal distance of 29 Å) has been reported to exhibit significant thermostabilization via the SpyRing system,^15^ the stabilization effect on TEVp appeared to be more modest. A recent study^26^ has proposed that interfacial interactions between SpyTag/SpyCatcher and the target enzyme may contribute to thermal stabilization, indicating that further optimization of these interfaces could enhance the stabilizing effect. To clarify these possibilities, future investigations involving not only structure prediction but also high-resolution structural analyses such as X-ray crystallography will be necessary.

Notably, all AlphaFold models constructed in this study included not only the enzyme domain but also a short peptide corresponding to the TEVp recognition site as a separate chain. This modeling strategy allowed us to verify that the substrate consistently occupied the correct position within the active site in each construct. Although this detail was not emphasized earlier in the manuscript, such confirmation is essential because even if SpyRing-mediated cyclization is successful, any misalignment of the substrate-binding site could compromise catalytic function. The fact that the substrate was properly accommodated in all models supports the conclusion that enzymatic activity was preserved in the cyclized constructs, as subsequently confirmed by experimental data.

Interestingly, this ability of AlphaFold 3 to model intermolecular interactions extends beyond enzyme-substrate complexes. In the case of SpyRing-mediated cyclization, SpyTag and SpyCatcher are fused within a single polypeptide chain, yet they must still adopt a favorable conformation to enable intramolecular complex formation. Our results demonstrate that AlphaFold accurately modeled this intramolecular association, despite the relatively long terminal distance of TEVp, indicating that such predictive capability can be generalized to the design of structurally constrained proteins. This feature represents a significant advancement in computational design, facilitating the rational engineering of proteins with enhanced stability or novel topologies.

To evaluate the potential utility of the enzymes under high-temperature conditions—such as those encountered in thermophilic organisms or industrial bioprocesses—we assessed the proteolytic activity of cTEVp at 50°C using the thermostable substrate ChBD. Under these conditions, cTEVp demonstrated significantly higher cleavage activity than both TEVp and ncTEVp (Figure 4A and 4B), as confirmed by quantitative analysis (*p* = 0.0049 and 0.00056, respectively), indicating its enhanced catalytic efficiency at elevated temperatures. Although previous studies have shown that SpyRing-mediated cyclization can promote refolding and recovery of enzyme activity after thermal denaturation,^13,15,27^ our results suggest that cyclization may also directly improve enzymatic function under continuous heat exposure. Given that our TEVp construct includes the same L56V and S135G mutations as TEVp^2M^,^7,8^ the superior activity of cTEVp at 50°C suggests that SpyRing cyclization further enhances stability beyond that conferred by these mutations alone. These findings extend the known benefits of SpyRing cyclization and highlight its applicability even for proteases that do not refold after heat stress, such as TEVp. Further biophysical analyses, including direct comparison with reversible enzymes such as β-lactamase, may help clarify the mechanistic basis of this effect.

Collectively, our results support the utility of SpyRing-mediated cyclization using discrete modular domains as a generalizable platform for enhancing protein stability. By integrating structure-based design via AlphaFold with genetically encodable cyclization domains, this strategy enables the rational engineering of robust and thermostable proteins with broad applicability in biocatalysis, therapeutic design, and synthetic biological devices. The successful cyclization of TEVp, despite its relatively long terminal separation, suggests that computational modeling can effectively guide the design of cyclized proteins, even in cases previously considered unsuitable for SpyRing. This work not only advances protein engineering but also exemplifies a synthetic biology framework for designing structurally constrained, thermostable biocatalysts for diverse industrial and biomedical applications.

## 4. Methods

### 4.1. Design and construction of expression plasmids

*E. coli* strain DH5α was used for plasmid preparation. *E. coli* cells were grown aerobically at 37 °C in Luria-Bertani (LB) medium (1% (w/v) polypeptone, 0.5% (w/v) yeast extract, and 0.5% (w/v) NaCl). For the construction of expression plasmids, restriction enzyme-digested fragments from existing plasmids, PCR products, or synthetic DNA were assembled using T4 DNA Ligase (Takara) or NEBuilder (New England Biolabs). Chemically synthesized oligonucleotides and double-stranded DNA fragments were purchased from Eurofins Genomics or GeneArt Strings (Thermo Fisher Scientific).

pHIT184 was constructed and used as an *E. coli* expression vector. This plasmid contains the expression cassette from pET-11a (Merck Millipore) and the replication origin from the pUC18 vector (GenBank: L08752.1) (Table S1, Figure S3). This plasmid retains the IPTG-inducible T7 promoter, the *lacO* operator and the *lacI* gene, which suppresses background expression, similar to the pET-11a vector. All plasmids used for protein production were subcloned into this vector backbone. Various target genes were subcloned into NdeI/BamHI or XbaI/BamHI site downstream of the T7 promoter to generate expression plasmids for the proteins listed in Table 1. The sequences of the constructed plasmids were confirmed by Sanger sequencing to ensure correct assembly.

The TEVp variant used throughout this study (L56V, S135G, S219V, Δ238–242), referred to as standard TEVp in the main text,^9^ has been widely employed due to its improved solubility and activity. Although its expression and purification have typically relied on the pDZ2087 plasmid,^9^ which requires co-expression of rare tRNA genes (e.g., from pRIL) due to biased codon usage, we circumvented this limitation by using a synthetic, codon-optimized TEVp gene for efficient expression in *E. coli*.

First, we constructed the plasmid pHIT184-MBP-TEVp for the expression of TEVp (Table S2). The amino acid sequence of the TEVp part in the recombinant fusion protein produced from this plasmid is identical to that of TEVp produced using pDZ2087,^9^ except for an additional 11 residues at the C-terminus, comprising a linker (ASA) and a Strep-tag (WSHPQFEK). The recombinant protein is produced in *E. coli* as a fusion protein with MBP derived from *E. coli* (Table S2), which promotes soluble expression.^25^ After expression, the TEVp undergoes *in vivo* self-cleavage at the TEVp cleavage site (TEVcs) located between MBP and TEVp,^9^ yielding active TEVp bearing an N-terminal polyhistidine tag (H_7_-tag) and a C-terminal Strep-tag.

For cyclized protein expression using the SpyRing system, the N-terminus and C-terminus of the target protein were fused with ST and SC, respectively. However, these fusions alone did not enhance soluble expression. Therefore, similar to the TEVp construct, cTEVp was expressed as a fusion protein with MBP, and the expression plasmid pHIT184-MBP-cTEVp was constructed (Table S2). In this study, to facilitate purification, all recombinant proteins were expressed with an N-terminal polyhistidine tag (H_6_- or H_7_-tag) and a C-terminal Strep tag. Additionally, we constructed an expression plasmid of non-cyclized TEVp (ncTEVp) (pHIT184-MBP-ncTEVp) incorporating the DA mutant,^28^ in which the aspartic acid residue in ST essential for isopeptide bond formation is substituted with alanine, to prepare a negative control that retains both ST and SC but cannot form the covalent bond (Table S2).

As substrates for TEVp, we used Mocr^21^ and T4 lysozyme (T4L),^22^ and the chitin-binding domain (ChBD)^23^ from *Thermococcus kodakarensis* KOD1. All three constructs contained a TEVcs between an N-terminal H_6_-tag and the substrate protein (Tables S2, S3).

### 4.2. Protein expression and purification

*E. coli* C41(DE3) cells harboring each expression plasmid (see Table S2) were cultured in a modified Studier autoinduction medium^29^ containing 2% polypeptone, 0.5% yeast extract, 0.5% NaCl, 0.6% Na_2_HPO_4_, 0.3% KH_2_PO_4_, 0.5% glycerol, 0.05% glucose, 0.2% lactose, supplemented with 100 μg/mL ampicillin. Cells were grown at 37 °C with shaking at 180 rpm until an optical density of ~0.1 was reached. At this point, the temperature was reduced according to the conditions optimized for each protein (see Table 1). The cultures were further incubated, allowing protein expression to be automatically induced upon glucose depletion and lactose metabolism. Cells were harvested by centrifugation at 6,000 g for 10 min and stored at −20 °C until use.

Frozen cell pellets (~3 g wet weight per 100 mL culture) were thawed, resuspended in phosphate-buffered saline (PBS; 10 mM sodium phosphate, 150 mM NaCl, pH 7.4), and supplemented with hen egg white lysozyme (final concentration: 1 mg/mL) and Triton X-100 (final concentration: 1% (v/v)). The suspension was incubated at 37 °C for 10 min to facilitate cell lysis. After incubation, a 2% (w/v) solution of streptomycin sulfate in PBS was added to a final concentration of 1% (w/v), and the mixture was gently inverted to ensure thorough mixing. The suspension was then placed on ice for 5 min to allow precipitation of nucleic acids. The lysate was subsequently centrifuged at 20,000 g for 30 min at 4 °C.

The resulting supernatant was loaded onto a Ni-affinity column packed with cOmplete His-Tag Purification Resin (Roche), pre-equilibrated with buffer A (25 mM Tris-HCl (pH 8.0), 500 mM NaCl, and 10% (w/v) glycerol). After washing the column with five bed volumes of buffer A, the bound protein was eluted using buffer A containing 500 mM imidazole. The eluted fractions were collected and loaded onto a Strep-Tactin Superflow High-Capacity Column (1 mL; IBA), pre-equilibrated with buffer A. The column was washed with 10 mL of buffer A, and the bound protein was eluted with buffer A containing 1 mM *d*-desthiobiotin. Fractions containing the target protein were pooled, aliquoted as needed, and stored at −80 °C. The purified protein was analyzed by SDS-PAGE using an Any kD denaturing polyacrylamide gel (Bio-Rad). The protein concentration was determined using the Bradford method with Protein Assay CBB Solution (5× concentrate) (Nacalai Tesque), using bovine serum albumin as the standard.

### 4.3. Protease activity assay

Protease activity was evaluated by assessing the cleavage of three substrates (Mocr, T4L, and ChBD), each containing a TEVp cleavage site downstream of the His tag. The activity of cTEVp was compared to that of the standard TEVp (without ST and SC) and the ncTEVp, which carries a mutation in the ST sequence preventing cyclization.

The cleavage reactions were performed using Mocr as the substrate. Enzyme and substrate were mixed in a reaction buffer containing 50 mM Tris-HCl (pH 8.0), 0.5 mM EDTA, and 1 mM dithiothreitol (DTT), and incubated at 37°C for 60 minutes. The substrate concentration was fixed at 100 μM, and enzyme-to-substrate molar ratios of 1:32, 1:128, and 1:512 were used. After incubation, 20 μL of the reaction mixture was collected, mixed with SDS-PAGE loading buffer containing 2% (v/v) β-mercaptoethanol, and heated at 100°C for 10 minutes to terminate the reaction. Cleavage activity was evaluated by SDS-PAGE using an Any kD SDS-PAGE gel (Bio-Rad), and the band intensities of the substrate and cleavage products were quantified using ImageJ software.^30^ For accurate quantification, images were scanned at 48-bit resolution, and the red channel was extracted and converted to 32-bit grayscale. Pixel values were log-transformed and scaled to approximate optical density, following a standard procedure involving logarithmic conversion (log_10_) after normalization. Integrated densities of each band were then measured after background subtraction.

For the assessment of residual soluble fraction and activity after heat treatment, His-tagged T4L, a substrate that is relatively resistant to cleavage, was used. This substrate was selected because, even when mixed with an equimolar amount of enzyme, 3 hours at 37°C are required to achieve 50% cleavage, making it suitable for simultaneous quantification of both the enzyme and substrate on SDS-PAGE gels. The assay was performed by incubating 20 μL of a 50 μM enzyme solution at 50°C for 0, 1, 2, 4, 8, 16, or 32 minutes. After incubation at 50°C, the enzyme solution was centrifuged at 15,000 × g for 10 minutes at 4°C. The resulting supernatant (10 μL) was then mixed with 10 μL of 50 μM T4L substrate, followed by incubation at 37°C for 3 hours before reaction termination. Protease activity was subsequently quantified using the method described above.

For additional evaluation of protease activity under moderate and elevated temperatures, comparative cleavage assays were conducted using T4 lysozyme (T4L) and the thermostable chitin-binding domain (ChBD) as substrates (Figure 4). For reactions with T4L, enzyme (25 μM, 10 μL) and substrate (100 μM, 10 μL) were mixed and incubated at either 30°C or 37°C for 3 hours under static conditions. After incubation, the reactions were terminated and analyzed by SDS-PAGE as described above. To achieve comparable cleavage levels to the T4L assay, a reduced amount of enzyme (25 μM, 2 μL) was used for reactions with ChBD (100 μM, 18 μL), and the reaction mixture was incubated for 30 minutes in a PCR thermal cycler block pre-set to 50°C. Reactions were terminated by heating the samples to 98°C for 2 minutes using the same thermal cycler, followed by standard sample processing as described above.

### 4.4. AlphaFold prediction and statistical analysis

Structural models of the various cTEVp constructs were generated using AlphaFold 3^19^ by inputting the amino acid sequences of cTEVp and its model substrate peptide (GENLYFQG) as two separate chains into the web-based prediction tool. Predictions were performed using the default settings with multimer mode enabled to allow modeling of intermolecular interactions. Although the interface allows specification of post-translational modifications (PTMs), the isopeptide bond formation between SpyTag and SpyCatcher is not currently supported; therefore, no PTMs were specified. Five ranked models were generated for each input, and the top-ranked model based on pLDDT was selected for further analysis. Structural models were manually inspected and adjusted using Coot, ^31^ and molecular graphics were rendered with PyMOL (Schrödinger, LLC).^32^

All experimental data are expressed as means ± standard deviations (SD) from at least two independent experiments. Statistical significance was determined using an unpaired two-tailed Student’s t-test, with *p* < 0.05 considered statistically significant. For figures with asterisks, significance levels are denoted as follows: \**p* < 0.05, \*\**p* < 0.01, \*\*\**p* < 0.001. All statistical analyses were performed using Microsoft Excel.

## Supporting information

Supplemental References

## Abbreviations

ChBD: chitin-binding domain
cTEVp: cyclized TEVp
ncTEVp: non-cyclized TEVp
SC: SpyCatcher
ST: SpyTag
T4L: T4 lysozyme
TEV: Tobacco Etch Virus
TEVcs: TEVp cleavage site
TEVp: TEV protease.

## Author Information

**Corresponding Author**

Tadashi Nakai – *Graduate School of Science and Technology, Hiroshima Institute of Technology, Hiroshima 731-5193, Japan; Faculty of Life Sciences, Hiroshima Institute of Technology, Hiroshima
731-5193, Japan*; orcid.org/0000-0001-9770-8168;

**Authors**

Yota Nakai – *Hiroshima University Senior High School, Hiroshima 734-0005, Japan*

Naoki Takami – *Faculty of Life Sciences, Hiroshima Institute of Technology, Hiroshima 731-5193, Japan*

Emi Nakai – *Faculty of Life Sciences, Hiroshima Institute of Technology, Hiroshima 731-5193, Japan*

Toshihide Okajima – *Institute of Scientific and Industrial Research (SANKEN), Osaka University, Ibaraki, Osaka 567-0047, Japan*

## Author Contributions

TN contributed to conceptualization, methodology, investigation, formal analysis, validation, writing—original draft preparation, writing—review and editing, project administration, supervision, and funding acquisition. YN contributed to methodology and investigation. NT and EN contributed to investigation. TO contributed to supervision, funding acquisition, and writing—review and editing. All authors read and approved the final version of the manuscript.

## Notes

The authors declare that they have no competing financial or non-financial interests.

## Acknowledgments

We thank Takashi Nagami, Ayumi Yamasaki, and Daisuke Kimura for their contributions to plasmid construction in the early stages of this work. The authors acknowledge the use of OpenAI’s ChatGPT for English language editing and text polishing during manuscript preparation. The authors reviewed and edited the content as needed and take full responsibility for the final version of the manuscript.

This work was supported by the Japan Society for the Promotion of Science (JSPS) KAKENHI Grant Numbers JP16K07691 and JP22K05419 awarded to T.N., and JP15K07391 and JP22K19150 awarded to T.O. This work was also supported by the Cooperative Research Program of the Network Joint Research Center for Materials and Devices.

## Notes

### Competing Interest Statement

The authors have declared no competing interest.

