## Supplementary material for "SpyRing-Mediated Cyclization of TEV Protease, Guided by AlphaFold, Improves Thermostability": Table S1

**Table S1. Nucleotide sequences used for the construction of the expression vector pHIT184.**

| | Nucleotide sequence[a] | Comment |
|---|---|---|
| 1 | CCCGGGtgcctaatgagtgagctaacttacattaattgcgttgcgctcactgcccgctttccagtcgggaaacc<br>tgtcgtgccagctgcattaatgaatcggccaacgcgcggggagaggcggtttgcgtattgggcgccagggtggt<br>ttttcttttcaccagtgagacgggcaacagctgattgcccttcaccgcctggccctgagagagttgcagcaagc<br>ggtccacgctggtttgccccagcaggcgaaaatcctgtttgatggtggttaacggcgggatataacatgagctg<br>tcttcggtatcgtcgtatcccactaccgagatGtccgcaccaacgcgcagcccggactcggtaatggcgcgcat<br>tgcgcccagcgccatctgatcgttggcaaccagcatcgcagtgggaacgatgccctcattcagcatttgcatgg<br>tttgttgaaaaccggacatggcactccagtcgccttcccgttccgctatcggctgaatttgattgcgagtgaga<br>tatttatgccagccagccagacgcagacgcgccgagacagaacttaatgggcccgctaacagcgcgatttgctg<br>gtgacccaatgcgaccagatgctccacgcccagtcgcgtaccgtcttcatgggagaaaataatactgttgatgg<br>gtgtctggtcagagacatcaagaaataacgccggaacattagtgcaggcagcttccacagcaatggcatcctgg<br>tcatccagcggatagttaatgatcagcccactgacgcgttgcgcgagaagattgtgcaccgccgctttacaggc<br>ttcgacgccgcttcgttctaccatcgacaccaccacgctggcacccagttgatcggcgcgagatttaatcgccg<br>cgacaatttgcgacggcgcgtgcagggccagactggaggtggcaacgccaatcagcaacgactgtttgcccgcc<br>agttgttgtgccacgcggttgggaatgtaattcagctccgccatcgccgcttccactttttcccgcgttttcgc<br>agaaacgtggctggcctggttcaccacgcgggaaacggtctgataagagacaccggcatactctgcgacatcgt<br>ataacgttactggtttcacattcaccaccctgaattgactctcttccgggcgctatcatgccataccgcgaaag<br>gttttgcgccattcgatggtgtccgggatctcgacgctctcccttatgcgactcctgcattaggaagcagccca<br>gtagtaggttgaggccgttgagcaccgccgccgcaaggaatggtgcatgcaaggagatggcgcccaacagtccc<br>ccggccacggggcctgccaccatacccacgccgaaacaagcgctcatgagcccgaagtggcgagcccgatcttc<br>cccatcggtgatgtcggcgatataggcgccagcaaccgcacctgtggcgccggtgatgccggccacgatgcgtc<br>cggcgtagaggatcgagatctcgatcccgcgaaattaatacgactcactatagggga attgtgagcggataaca<br>attcccctctagaaataattttgtttaactttaagaaggagatatacatatgatgaaacatctgacgcccgcca<br>atgccaaggcaaaggcctttgtcgaagccgaggccgagggccgcgaggaggaggtcgtggcgatgaatgcgctg<br>gtcggctgcgctagccatcaccatcaccatcactaggatccggctgctaacaaagcccgaaaggaagctgagtt<br>ggctgctgccaccgctgagcaataactagcataaccccttggggcctctaaacgggtcttgaggggttttttgc<br>tgaaaggaggaactatatccggatGGGAAGGCCT | XmaI - *lacI* - T7 promoter - XbaI - NdeI - a short ORF - BamHI – T7 terminator - StuI [b]<br><br>'gatGtcc' represents an EcoRV site that was removed by site-directed mutagenesis. |
| 2 | AGGCCTTtaccaatgcttaatcagtgaggcacctatctcagcgatctgtctatttcgttcatccatagttgcctg<br>actccccgtcgtgtagataactacgatacgggagggcttaccatctggccccagtgctgcaatgataccgcgag<br>aGccacgctcaccggctccagatttatcagcaataaaccagccagccggaagggccgagcgcagaagtggtcct<br>gcaactttatccgcctccatccagtctattaattgttgccgggaagctagagtaagtagttcgccagttaatag<br>tttgcgcaacgttgttgccattgctacaggcatcgtggtgtcacgctcgtcgtttggtatggcttcattcagct<br>ccggttcccaacgatcaaggcgagttacatgatcccccatgttgtgcaaaaaagcggttagctccttcggtcct<br>ccgatcgttgtcagaagtaagttggccgcagtgttatcactcatggttatggcagcactgcataattctcttac<br>tgtcatgccatccgtaagatgcttttctgtgactggtgagtactcaaccaagtcattctgagaatagtgtatgc<br>ggcgaccgagttgctcttgcccggcgtcaatacgggataataccgcgccacatagcagaactttaaaagtgctc<br>atcattggaaaacgttcttcggggcgaaaactctcaaggatcttaccgctgttgagatccagttcgatgtaacc<br>cactcgtgcacccaactgatcttcagcatcttttactttcaccagcgtttctgggtgagcaaaaacaggaaggc<br>aaaatgccgcaaaaaagggaataagggcgacacggaaatgttgaatactcatactcttcctttttcaatattat<br>tgaagcatttatcagggttattgtctcatgagcggatacatatttgaatgtatttagaaaaataaacaaatagg<br>gGCATGC | StuI - Ap[R] - SphI [c]<br><br>'gagaGc' represents a BsaI site that was removed by site-directed mutagenesis. |
| 3 | GCATGCgacccttgtcttttccgctgcataaccctgcttcggggtcattatagcgatttttcggtatatcca<br>tcctttttcgcacgatatacaggattttgccaaagggttcgtgtagactttccttggtgtatccaacggcgtca<br>gccgggcaggataggtgaagtaggcccacccgcgagcgggtgttccttcttcactgtcccttattcgcacctgg<br>cggtgctcaacgggaatcctgctctgcgaggctggccgtaggccggccgcgatgcaggtggctgctgaacccc<br>agccggaactgaccccacaaggccCTGCAG | SphI - *oriT* - PstI [d] |
| 4 | CTGCAGtcatgaccaaaatcccttaacgtgagttttcgttccactgagcgtcagaccccgtagaaaagatcaaa<br>ggatcttcttgagatcctttttttctgcgcgtaatctgctgcttgcaaacaaaaaaaccaccgctaccagcggt<br>ggtttgtttgccggatcaagagctaccaactctttttccgaaggtaactggcttcagcagagcgcagataccaa<br>atactgttcttctagtgtagccgtagttaggccaccacttcaagaactctgtagcaccgcctacatacctcgct<br>ctgctaatcctgttaccagtggctgctgccagtggcgataagtcgtgtcttaccgggttggactcaagacgata<br>gttaccggataaggcgcagcggtcgggctgaacggggggttcgtgcacacagcccagcttggagcgaacgacct<br>acaccgaactgagatacctacagcgtgagctatgagaaagcgccacgcttcccgaagggagaaaggcggacagg<br>tatccggtaagcggcagggtcggaacaggagagcgcacgagggagcttccagggggaaacgcctggtatcttta<br>tagtcctgtcgggtttcgccacctctgacttgagcgtcgatttttgtgatgctcgtcaggggggcggagcctat<br>ggaaaaacgccagcaacgcggcctttttacggttcctggccttttgctggccttttgctCCCGGG | PstI - ColE1 origin - XmaI [e] |

The pHIT184 plasmid was constructed by assembling Fragments 1–4 listed in this table.
[a] For fragment sequences, lowercase letters indicate nucleotide sequences identical to the template in PCR-amplified fragments or to published sequences for chemically synthesized fragments, while uppercase letters represent modifications or additional nucleotides introduced in this study. In practice, additional nucleotides were included at both termini to facilitate efficient digestion and assembly.

[b] Fragment 1 was obtained by PCR using pET-11a derivative, pET-QhpC$_{(-28)-7}$-H6.[S1] The PCR product was amplified from this plasmid template, as the original empty pET-11a vector was unavailable. Instead, a derivative containing a short ORF, QhpC$_{(-28)-7}$-H6, was used.
[c] Fragment 2 was obtained by PCR using pUC18 (GenBank: L08752.1) as the template.
[d] Fragment 3 corresponds to the nucleotide sequence of pSEVA121 (GenBank: JX560322.1). Instead of PCR amplification, a synthetic DNA fragment matching this sequence was used. This region contains *oriT*, which is useful for conjugation-based plasmid transfer in other studies. However, *oriT* was not utilized in this study as conjugation was not performed.
[e] Fragment 4 was obtained by PCR using pUC18 (GenBank: L08752.1) as the template.
