## Supplementary material for "SpyRing-Mediated Cyclization of TEV Protease, Guided by AlphaFold, Improves Thermostability": Table S2

**Table S2. Nucleotide sequences used for the construction of expression plasmids.**

| | Nucleotide sequence [a] | Comment |
|---|---|---|
| 1 | CATATGaaaatcgaagaaggtaaactggtaatctggattaacggcgataaaggctataacggtctcgctgaag<br>tcggtaagaaattcgagaaagataccggaattaaagtcaccgttgagcatccggataaactggaagagaaatt<br>cccacaggttgcggcaactggcgatggccctgacattatcttctgggcacacgaccgctttggtggctacgct<br>caatctggcctgttggctgaaatcaccccggacaaagcgttccaggacaagctgtatccgtttacctgggatg<br>ccgtacgttacaacggcaagctgattgcttacccgatcgctgttgaagcgttatcgctgatttataacaaaga<br>tctgctgccgaacccgccaaaaacctgggaagagatcccggcgctggataaagaactgaaagcgaaaggtaag<br>agcgcgctgatgttcaacctgcaagaaccgtacttcacctggccgctgattgctgctgacggggttatgcgt<br>tcaagtatgaaaacggcaagtacgacattaaagacgtgggcgtggataacgctggcgcgaaagcgggtctgac<br>cttcctggttgacctgattaaaaacaaacacatgaatgcagacaccgattactccatcgcagaagctgccttt<br>aataaaggcgaaacagcgatgaccatcaacggcccgtgggcatggtccaacatcgacaccagcaaagtgaatt<br>atggtgtaacggtactgccgaccttcaagggtcaaccatccaaaccgttcgttggcgtgctgagcgcaggtat<br>taacgccgccagtccgaacaaagagctggcgaaagagttcctcgaaaactatctgctgactgatgaaggtctg<br>gaagcggttaataaagacaaaccgctgggtgccgtagcgctgaagtcttacgaggaagagttggcgaaagatc<br>cacgtattgccgccaccatggaaaacgcccagaaaggtgaaatcatgccgaacatcccgcagatgtccgcttt<br>ctggtatgccgtgcgtactgcggtgatcaacgccgccagcggtcgtcagactgtcgatgaagccctgaaagac<br>gcgcagactaatGCGGCCGC | NdeI - MBP - NotI [b] |
| 2 | GCGGCCGCAaacaataacaataataacaacaacaataatctgggaattgaaggtcgcggtgagaacctgtact<br>tccaagggcatcatcaccatcatcatcatggcgaatccctgtttaaaggcccacgtgactataacccgatttc<br>ctccaccatttgccacctgaccaatgaaagcgatggccatacaacctcactgtatgggattggttttggcccc<br>tttatcatcacgaacaaacacctgtttcgccgcaacaatggtactttggtggtacagtcgttacatggcgtgt<br>tcaaagtcaagaatacgacgacattacagcagcatctgatcgatggacgcgacatgattatcatccgtatgcc<br>caaagattttccgccttttccacagaaactcaagtttcgcgaaccgcaacgtgaggaacggatttgccttgta<br>acgaccaactttcaaaccaaaagtatgagctcaatggtttcggatacgagctgtaccttcccaagcggtgatg<br>gcatcttctggaaacactggattcagacaaaagacggtcaatgtggcagtccgttagtgtctactcgcgatgg<br>cttcattgtgggcattcactcggcaagcaatttcaccaacacgaataactacttcacttctgtgcctaagaac<br>tttatggagcttctgaccaatcaggaagcccagcagtgggttagtggttggcgtctgaatgcggacagcgttc<br>tctggggtgggcacaaagtgttcatggtcaaaccggaagaaccgtttcaaccggtcaaagaggctacccagtt<br>gatgaacgctagcgcttggagccacccgcagttcgaaaaataGGATCC | NotI - Factor Xa cleavage site - TEVcs - $H_7$-tag - TEVp - NheI - Strep-tag - BamHI [c] |
| 3 | GCGGCCGCAaacaataacaataataacaacaacaataatctgggaattgaaggtcgcggtgagaacctgtact<br>tccaagggcatcatcaccatcatcatCATATG | NotI - N-tag - TEVcs - NdeI [d] |
| 4 | CATATGtccgcgcatattgtcatggttgatgcgtacaaaccgaccaaaGCGGCCGC | NdeI - ST - NotI [e] |
| 4a | CATATGtccgcgcatattgtcatggttgCtgcgtacaaaccgaccaaaGCGGCCGC | NdeI – STmut - NotI [f] |
| 5 | GCGGCCGCactgtttaaaggcccacgtgactataacccgatttcctccaccatttgccacctgaccaatgaaa<br>gcgatggccatacaacctcactgtatgggattggttttggcccctttatcatcacgaacaaacacctgtttcg<br>ccgcaacaatggtactttggtggtacagtcgttacatggcgtgttcaaagtcaagaatacgacgacattacag<br>cagcatctgatcgatggacgcgacatgattatcatccgtatgcccaaagattttccgccttttccacagaaac<br>tcaagtttcgcgaaccgcaacgtgaggaacggatttgccttgtaacgaccaactttcaaaccaaaagtatgag<br>ctcaatggtttcggatacgagctgtaccttcccaagcggtgatggcatcttctggaaacactggattcagaca<br>aaagacggtcaatgtggcagtccgttagtgtctactcgcgatggcttcattgtgggcattcactcggcaagca<br>atttcaccaacacgaataactacttcacttctgtgcctaagaactttatggagcttctgaccaatcaggaagc<br>ccagcagtgggttagtggttggcgtctgaatgcggacagcgttctctggggtgggcacaaagtgttcatggtc<br>aaaccggaagaaccgtttcaaccggtcaaagaggctacccagttgatgaacGCTAGC | NotI- TEVp- NheI [g] |
| 6 | GCTAGCagcgattctgcgactcacatcaaattttccaaacgtgatgaagatggtaaggaactggctggcgcga<br>cgatggaacttcgggattcttccggcaaaacaattagcacgtggattagtgatggccaagtcaaagacttcta<br>tctgtatccggggaaatacacctttgtcgaaacagccgcaccagacggctatgaagtcgcgaccgcaatcacc<br>ttcacagtgaatgaacagggtcaggttaccgtgaatggaggcgcttggagccacccgcagttcgaaaaataGG<br>ATCC | NheI - SC - Strep-tag - BamHI [h] |
| 7 | TCTAGAaataattttgtttaactttaagaaggagatataccatgggcagcagccatcatcatcatcatcacga<br>ttacgatatcccaacgaccgaaaacctttacttccagggcGCGGCCGC | XbaI - $H_6$-tag - TEVcs - NotI [i] |
| 8 | GCGGCCGCAagcaatatgacctataataacgtttttgatcatgcctatgaaatgctgaaagaaaacatccgct<br>atgatgatattcgcgataccgatgatctgcatgatgcaattcacatggcagcagataatgcagttccgcatta<br>ttatgcagatattcgtagcgttatggccagcgaaggtattgatctggaatttgaagatagcggtctgatgccg<br>gataccaaagatgacattcgtattctgcaagcccgtatttatgaacagctgacaattgatctgtgggaagatg<br>cagaggatctgctgaatgaatatctggaagaggtggaagaatatgaagaggacgaagagGGTGGCAGCGGTGG<br>TGCTAGC | NotI - Mocr - GGSGG linker - NheI [j] |

| | | |
|---|---|---|
| 9 | GCTAGCgcttggagccacccgcagttcgaaaaataGGATCC | NheI - Strep-tag - BamHI [k] |
| 10 | TCTAGAaataattttgtttaactttaagaaggagatataccatgggcagcagccatcatcatcatcatcacgattacgatatcccaacgaccgaaaacctttacttccagggcCATATG | XbaI - $H_6$-tag - TEVcs - NdeI [l] |
| 11 | CATATGaatatatttgaaatgttacgtatagatgaaggtcttagacttaaaatctataaagacacagaaggctattacactattggcatcggtcatttgcttacaaaaagtccatcacttaatgctgctaaatctgaattagataaagctattgggcgtaatactaatggtgtaattacaaaagatgaggctgaaaaactctttaatcaggatgttgatgctgctgttcgcggcattctgagaaatgctaaattaaaaccggtttatgattctcttgatgcggttcgtcgcgctgcattgattaatatggttttccaaatgggagaaaccggtgtggcaggatttactaactctttacgtatgcttcaacaaaaacgctgggatgaagcagcagttaacttagctaaaagtagatggtataatcaaacacctaatcgcgcaaaacgagtcattacaacgtttagaactggcacttgggacgcgtatACTAGT | NdeI - T4L - NheI [m] |
| 12 | CATATGggcgactttgttaaaccgggtagcctgagcgttaaagttaccgattggggtaataccgaatatgacgttaccctgaatttaggtggcacctatgactgggttgttaaagtgaaactgaaagatggtagcagcgttagcagcttttggagcgcaaataaagccgaagaaggtggttacgttgtttttacaccggttagctggaatcgtggtccgaccgcaacctttggttttattgcaaccggtagcgaaagcgttgaagccatttatctgtatgttgatggtcagctgtgggatgcatggccgagcaatacccagcagccggaagagGGTGGTACTAGT | NdeI - ChBD - GG linker - SpeI [n] |

- The pHIT184-MBP-TEVp plasmid was constructed by assembling Fragments 1 and 2 into the NdeI/BamHI site of pHIT184.
- The pHIT184-MBP-cTEVp plasmid was constructed by assembling Fragments 1, 3, 4, 5, and 6 into the NdeI/BamHI site of pHIT184. The mutant plasmid pHIT184-MBP-ncTEVp carries the sequence that would result from using Fragment 4a instead of Fragment 4; however, the mutation was introduced directly into the plasmid by PCR-based site-directed mutagenesis.
- The pHIT184-Mocr plasmid was constructed by assembling Fragments 7, 8, 9 into the XbaI/BamHI site of pHIT184.
- The pHIT184-T4L plasmid was constructed by assembling Fragments 10 and 11 into the XbaI/NheI site of pHIT184.
- The pHIT184-ChBD plasmid was constructed by assembling Fragments 10 and 12 into the XbaI/NheI site of pHIT184.

[a] For fragment sequences, lowercase letters indicate nucleotide sequences identical to the template in PCR-amplified fragments or to published sequences for chemically synthesized fragments. When nucleotide sequences were derived from published amino acid sequences, they were also represented in lowercase. Uppercase letters represent modifications or additional nucleotides introduced in this study. In practice, additional nucleotides were included at both termini to facilitate efficient digestion and assembly.
[b] Fragment 1 was obtained by PCR using a colony of *E. coli* C41(DE3) as the template.
[c] Fragment 2 was chemically synthesized based on the amino acid sequence of TEVp and its upstream region from plasmid pDZ2087.[9]
[d] Fragment 3 was obtained by PCR using Fragment 2 as the template.
[e] Fragment 4 was chemically synthesized based on the amino acid sequence of SpyTag.[12]
[f] The mutation shown in Fragment 4a was obtained by PCR-based site-directed mutagenesis.
[g] Fragment 5 was obtained by PCR using Fragment 2 as the template.
[h] Fragment 6 was chemically synthesized based on the amino acid sequence of SC.[12]
[i] Fragment 7 was obtained by PCR using pET-His6-TEV-QhpG[S2] as the template.
[j] Fragment 8 was chemically synthesized based on the amino acid sequence of Mocr.[21]
[k] Fragment 9 was chemically synthesized based on the amino acid sequence of Strep-tag.[S3.]
[l] Fragment 10 was obtained by PCR using pET-His6-TEV-QhpG[S2] as the template.
[m] Fragment 11 was chemically synthesized based on the amino acid sequence of T4 lysozyme (T4L) .[22]
[n] Fragment 12 was chemically synthesized based on the amino acid sequence of the second chitin binding domain (ChBD)[23] of chitinase from *Thermococcus kodakarensis* KOD1 (GenBank: BAA88380.1).
