## Supplementary material for "SpyRing-Mediated Cyclization of TEV Protease, Guided by AlphaFold, Improves Thermostability": Table S3

**Table S3. Expressed amino acid sequences.**

| Name | Amino acid sequence [a] | Comment [a] |
|---|---|---|
| MBP-TEVp [b] | MKIEEGKLVIWINGDKGYNGLAEVGKKFEKDTGIKVTVEHPDKLEEKFPQVAATGDGPDIIFWAH<br>DRFGGYAQSGLLAEITPDKAFQDKLYPFTWDAVRYNGKLIAYPIAVEALSLIYNKDLLPNPPKTW<br>EEIPALDKELKAKGKSALMFNLQEPYFTWPLIAADGGYAFKYENGKYDIKDVGVDNAGAKAGLTF<br>LVDLIKNKHMNADTDYSIAEAAFNKGETAMTINGPWAWSNIDTSKVNYGVTVLPTFKGQPSKPFV<br>GVLSAGINAASPNKELAKEFLENYLLTDEGLEAVNKDKPLGAVALKSYEEELAKDPRIAATMENA<br>QKGEIMPNIPQMSAFWYAVRTAVINAASGRQTVDEALKDAQTNAAANNNNNNNNNNLGIEGRGEN<br>LYFQ/GHHHHHHHGESLFKGPRDYNPISSTICHLTNESDGHTTSLYGIGFGPFIITNKHLFRRNN<br>GTLVVQSLHGVFKVKNTTTLQQHLIDGRDMIIIRMPKDFPPFPQKLKFREPQREERICLVTTNFQ<br>TKSMSSMVSDTSCTFPSGDGIFWKHWIQTKDGQCGSPLVSTRDGFIVGIHSASNFTNTNNYFTSV<br>PKNFMELLTNQEAQQWVSGWRLNADSVLWGGHKVFMVKPEEPFQPVKEATQLMNASAWSHPQFEK<br>* | MBP - NotI - Factor Xa cleavage site - TEVcs - $H_7$-tag - TEVp - NheI - Strep-tag |
| MBP-cTEVp | MKIEEGKLVIWINGDKGYNGLAEVGKKFEKDTGIKVTVEHPDKLEEKFPQVAATGDGPDIIFWAH<br>DRFGGYAQSGLLAEITPDKAFQDKLYPFTWDAVRYNGKLIAYPIAVEALSLIYNKDLLPNPPKTW<br>EEIPALDKELKAKGKSALMFNLQEPYFTWPLIAADGGYAFKYENGKYDIKDVGVDNAGAKAGLTF<br>LVDLIKNKHMNADTDYSIAEAAFNKGETAMTINGPWAWSNIDTSKVNYGVTVLPTFKGQPSKPFV<br>GVLSAGINAASPNKELAKEFLENYLLTDEGLEAVNKDKPLGAVALKSYEEELAKDPRIAATMENA<br>QKGEIMPNIPQMSAFWYAVRTAVINAASGRQTVDEALKDAQTNAAANNNNNNNNNNLGIEGRGEN<br>LYFQ/GHHHHHHHMSAHIVMVDAYKPTKAAALFKGPRDYNPISSTICHLTNESDGHTTSLYGIGF<br>GPFIITNKHLFRRNNGTLVVQSLHGVFKVKNTTTLQQHLIDGRDMIIIRMPKDFPPFPQKLKFRE<br>PQREERICLVTTNFQTKSMSSMVSDTSCTFPSGDGIFWKHWIQTKDGQCGSPLVSTRDGFIVGIH<br>SASNFTNTNNYFTSVPKNFMELLTNQEAQQWVSGWRLNADSVLWGGHKVFMVKPEEPFQPVKEAT<br>QLMNASSDSATHIKFSKRDEDGKELAGATMELRDSSGKTISTWISDGQVKDFYLYPGKYTFVETA<br>APDGYEVATAITFTVNEQGQVTVNGGAWSHPQFEK* | MBP - NotI - Factor Xa cleavage site - TEVcs - $H_7$-tag - NdeI - ST - NotI - TEVp - NheI - SC - Strep-tag |
| MBP-ncTEVp | MKIEEGKLVIWINGDKGYNGLAEVGKKFEKDTGIKVTVEHPDKLEEKFPQVAATGDGPDIIFWAH<br>DRFGGYAQSGLLAEITPDKAFQDKLYPFTWDAVRYNGKLIAYPIAVEALSLIYNKDLLPNPPKTW<br>EEIPALDKELKAKGKSALMFNLQEPYFTWPLIAADGGYAFKYENGKYDIKDVGVDNAGAKAGLTF<br>LVDLIKNKHMNADTDYSIAEAAFNKGETAMTINGPWAWSNIDTSKVNYGVTVLPTFKGQPSKPFV<br>GVLSAGINAASPNKELAKEFLENYLLTDEGLEAVNKDKPLGAVALKSYEEELAKDPRIAATMENA<br>QKGEIMPNIPQMSAFWYAVRTAVINAASGRQTVDEALKDAQTNAAANNNNNNNNNNLGIEGRGEN<br>LYFQ/GHHHHHHHMSAHIVMVAAYKPTKAAALFKGPRDYNPISSTICHLTNESDGHTTSLYGIGF<br>GPFIITNKHLFRRNNGTLVVQSLHGVFKVKNTTTLQQHLIDGRDMIIIRMPKDFPPFPQKLKFRE<br>PQREERICLVTTNFQTKSMSSMVSDTSCTFPSGDGIFWKHWIQTKDGQCGSPLVSTRDGFIVGIH<br>SASNFTNTNNYFTSVPKNFMELLTNQEAQQWVSGWRLNADSVLWGGHKVFMVKPEEPFQPVKEAT<br>QLMNASSDSATHIKFSKRDEDGKELAGATMELRDSSGKTISTWISDGQVKDFYLYPGKYTFVETA<br>APDGYEVATAITFTVNEQGQVTVNGGAWSHPQFEK* | MBP - NotI - Factor Xa cleavage site - TEVcs - $H_7$-tag - NdeI – STmut - NotI - TEVp - NheI - SC - Strep-tag |
| Mocr | MGSSHHHHHHDYDIPTTENLYFQ/GAAASNMTYNNVFDHAYEMLKENIRYDDIRDTDDLHDAIHM<br>AADNAVPHYYADIRSVMASEGIDLEFEDSGLMPDTKDDIRILQARIYEQLTIDLWEDAEDLLNEY<br>LEEVEEYEEDEEGGSGGASAWSHPQFEK* | $H_6$-tag - TEVcs - NotI - Mocr - NheI - Strep-tag |
| T4L | MGSSHHHHHHDYDIPTTENLYFQ/GHMNIFEMLRIDEGLRLKIYKDTEGYYTIGIGHLLTKSPSL<br>NAAKSELDKAIGRNTNGVITKDEAEKLFNQDVDAAVRGILRNAKLKPVYDSLDAVRRAALINMVF<br>QMGETGVAGFTNSLRMLQQKRWDEAAVNLAKSRWYNQTPNRAKRVITTFRTGTWDAYASAWSHPQ<br>FEK* | $H_6$-tag - TEVcs - NdeI - T4L - NheI - Strep-tag |
| ChBD | MGSSHHHHHHDYDIPTTENLYFQ/GHMGDFVKPGSLSVKVTDWGNTEYDVTLNLGGTYDWVVKVK<br>LKDGSSVSSFWSANKAEEGGYVVFTPVSWNRGPTATFGFIATGSESVEAIYLYVDGQLWDAWPSN<br>TQQPEEGGTSAWSHPQFEK* | $H_6$-tag - TEVcs - NdeI - ChBD - SpeI/NheI - Strep-tag |

[a] ‘/’ denotes the cleavage position within the TEV recognition sequence.
[b] The differences between the amino acid sequence translated from the nucleotide sequence of pDZ2087[9] and that used in this study are found at three positions: the N-terminal ‘MKIEE’ (‘MKTEE’ in pDZ2087), ‘NAAAN’ (‘NSSSN’ in pDZ2087) containing the NotI site, and the C-terminal ‘QLMNASAWSHPQFEK*’ (‘QLMN*’ in pDZ2087).
