## Supplementary material for "SpyRing-Mediated Cyclization of TEV Protease, Guided by AlphaFold, Improves Thermostability": Figure S1

**A**

cTEVp-ver.1

HMSAHIVMVDAYKPTKGSGGSGLFKGPRDYNPISSTICHLTNESDGHTTSL
YGIGFGPFIITNKHLFRRNNGTLVVQSLHGVFKVKNTTTLQQHLIDGRDMI
IIRMPKDFPPFPQKLKFREPQREERICLVTTNFQTKSMSSMVSDTSCTFPS
GDGIFWKHWIQTKDGQCGSPLVSTRDGFIVGIHSASNFTNTNNYFTSVPKN
FMELLTNQEAQQWVSGWRLNADSVLWGGHKVFMVKPEEPFQPVKEATQLMN
GSGGSGGAMVDTLSGLSSEQGQSGDMTIEEDSATHIKFSKRDEDGKELAGA
TMELRDSSGKTISTWISDGQVKDFYLYPGKYTFVETAAPDGYEVATAITFT
VNEQGQVTVNG

**B**

cTEVp-ver.2

HMSAHIVMVDAYKPTKAAA---LFKGPRDYNPISSTICHLTNESDGHTTSL
YGIGFGPFIITNKHLFRRNNGTLVVQSLHGVFKVKNTTTLQQHLIDGRDMI
IIRMPKDFPPFPQKLKFREPQREERICLVTTNFQTKSMSSMVSDTSCTFPS
GDGIFWKHWIQTKDGQCGSPLVSTRDGFIVGIHSASNFTNTNNYFTSVPKN
FMELLTNQEAQQWVSGWRLNADSVLWGGHKVFMVKPEEPFQPVKEATQLMN
ASS----------------------------DSATHIKFSKRDEDGKELAGA
TMELRDSSGKTISTWISDGQVKDFYLYPGKYTFVETAAPDGYEVATAITFT
VNEQGQVTVNG

**C**

cTEVp-ver.3

HMSAHIVMVDAYKPT-AA------------YNPISSTICHLTNESDGHTTSL
YGIGFGPFIITNKHLFRRNNGTLVVQSLHGVFKVKNTTTLQQHLIDGRDMI
IIRMPKDFPPFPQKLKFREPQREERICLVTTNFQTKSMSSMVSDTSCTFPS
GDGIFWKHWIQTKDGQCGSPLVSTRDGFIVGIHSASNFTNTNNYFTSVPKN
FMELLTNQEAQQWVSGWRLNADSVLWGGHKVFMVKPPFPKETLN--------
ASS-----------------------------SATHIKFSKRDEDGKELAGA
TMELRDSSGKTISTWISDGQVKDFYLYPGKYTFVETAAPDGYEVATAITFT
VNEQGQVTVNG

**D**

cTEVp-ver.4

HMSAHIVMVDAYKPT-AA------------NPISSTICHLTNESDGHTTSL
YGIGFGPFIITNKHLFRRNNGTLVVQSLHGVFKVKNTTTLQQHLIDGRDMI
IIRMPKDFPPFPQKLKFREPQREERICLVTTNFQTKSMSSMVSDTSCTFPS
GDGIFWKHWIQTKDGQCGSPLVSTRDGFIVGIHSASNFTNTNNYFTSVPKN
FMELLTNQEAQQWVSGWRLNADSVLWGGHKVFMVKPPFPKETLN--------
ASS-----------------------------SATHIKFSKRDEDGKELAGA
TMELRDSSGKTISTWISDGQVKDFYLYPGKYTFVETAAPDGYEVATAITFT
VNEQGQVTVNG

**Figure S1. Amino acid sequences used for AlphaFold structure prediction of cTEVp variants.** Amino acid sequences corresponding to the cTEVp designs shown in Figure 1A–D were used as input for AlphaFold structural predictions. Residues are color-coded to match the domain and linker annotations in Figure 1: N-terminal SpyTag (yellow), flexible linker (magenta), C-terminal SpyCatcher (green), TEVp catalytic domain (cyan). The sequence shown for cTEVp-ver.2 is identical to that listed in Table S3; it is included here again to allow visual comparison to other variants. In cTEVp-ver.4, a single amino acid residue (Tyr) was deleted from the linker region relative to cTEVp-ver.3, as indicated by a white dash (–) on a red background, marking the deletion site in the aligned sequence.
