## Supplementary material for "SpyRing-Mediated Cyclization of TEV Protease, Guided by AlphaFold, Improves Thermostability": Figure S2

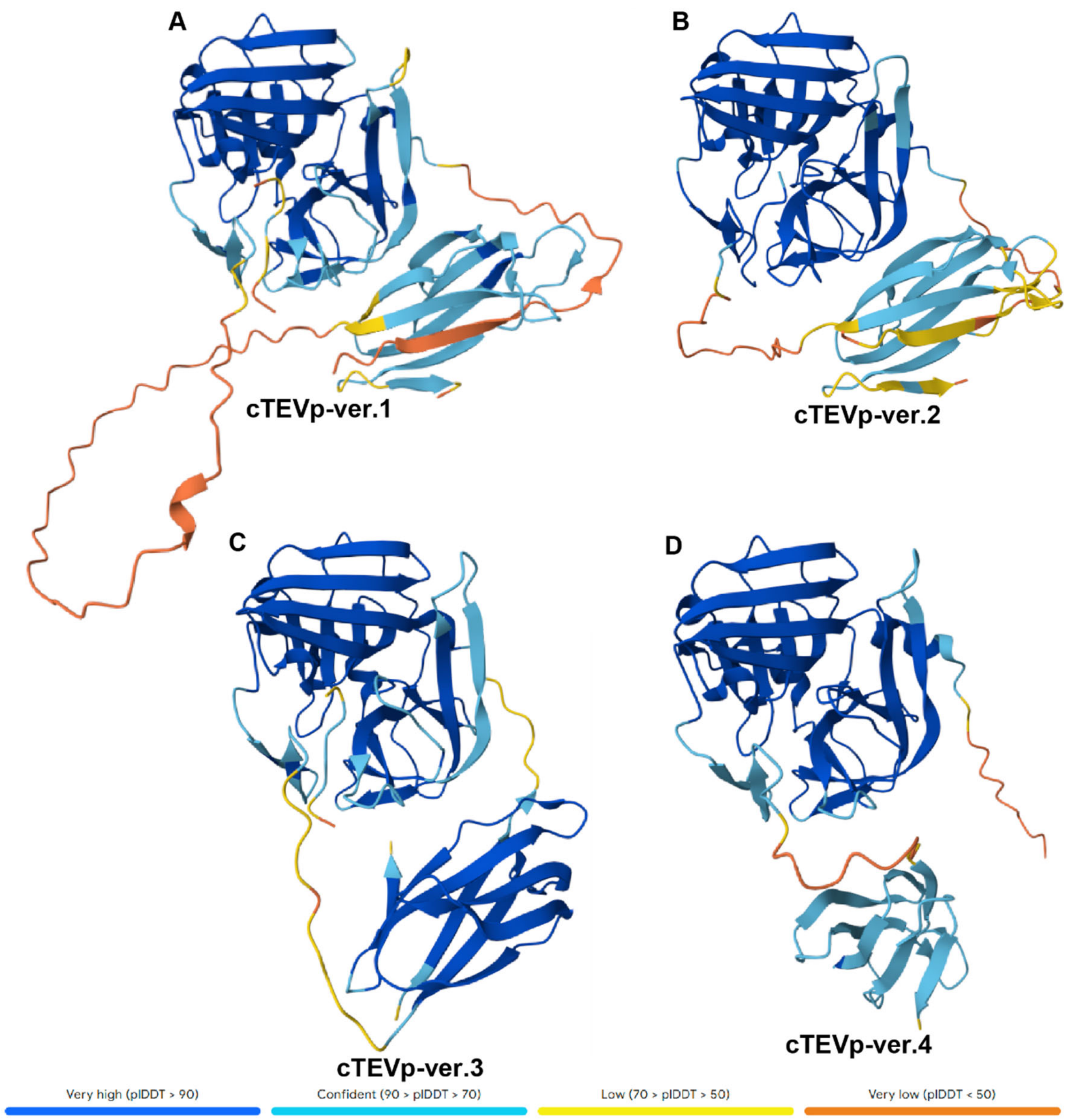


**Figure S2. AlphaFold-predicted structures of cTEVp variants colored by per-residue confidence scores (pLDDT).**

Predicted structures of cTEVp variants (ver.1–ver.4), corresponding to those shown in Figure 1A–D, are displayed with color-coded per-residue pLDDT scores as predicted by AlphaFold. The models are oriented to match those in Figure 1. Residue confidence levels are indicated as follows: very high (pLDDT > 90, dark blue), confident (90 > pLDDT > 70, light blue), low (70 > pLDDT > 50, yellow), and very low (pLDDT < 50, orange).
