## Supplementary material for "SpyRing-Mediated Cyclization of TEV Protease, Guided by AlphaFold, Improves Thermostability": Figure S3

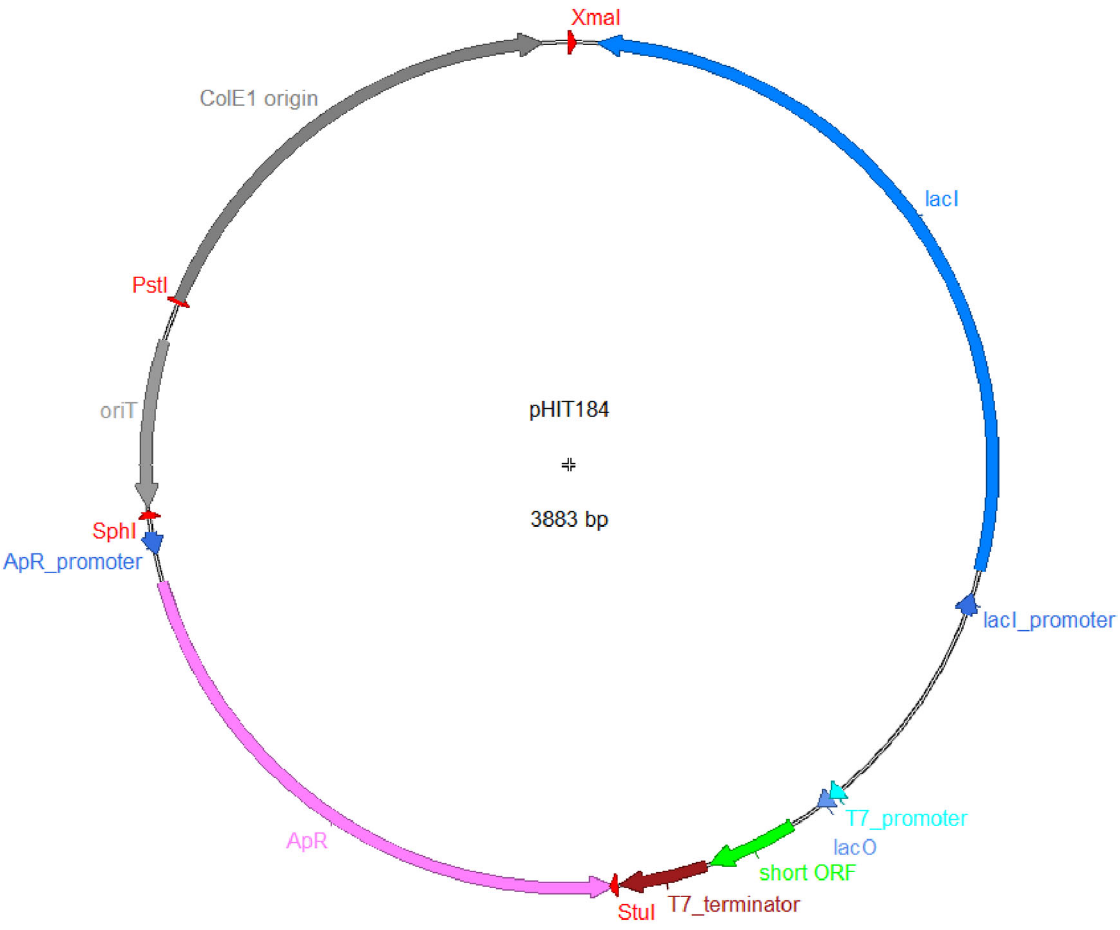


**Figure S3. Plasmid map of pHIT184.**

pHIT184 is an *E. coli* expression vector designed for high-level protein production using the T7 promoter system. The plasmid comprises a pUC-derived ColE1 origin of replication, an ampicillin resistance (ApR) gene driven by its native promoter, and an expression cassette derived from the pET series (Merck Millipore). This cassette includes the *lacI* gene under the control of the *lacI* promoter, allowing tight repression of expression, and a T7 promoter regulated by the *lac* operator (*lacO*) sequence. A short open reading frame (ORF) is present downstream of the T7 promoter to allow insertion of a gene of interest. The T7 transcription unit is terminated by a T7 terminator. Restriction enzyme sites that appear in Table S1 are indicated in the map by red arrowheads. The map was generated using the ApE (A plasmid Editor) software.[S4]
